# Maternal inflammation has a profound effect on cortical interneuron development in a stage and subtype-specific manner

**DOI:** 10.1101/412981

**Authors:** Navneet A. Vasistha, Maria Pardo-Navarro, Janina Gasthaus, Dilys Weijers, Michaela K. Müller, Diego García-González, Susmita Malwade, Irina Korshunova, Ulrich Pfisterer, Jakob von Engelhardt, Karin S. Hougaard, Konstantin Khodosevich

## Abstract

Severe infections during pregnancy are one of the major risk factors for cognitive brain impairment in offspring. It has been suggested that maternal inflammation leads to dysfunction of cortical GABAergic interneurons that in turn underlies cognitive impairment of the affected offspring. However, the evidence comes largely from studies of adult or mature brain and how impairment of inhibitory circuits arises upon maternal inflammation is unknown. Here we show that maternal inflammation affects multiple steps of cortical GABAergic interneuron development, i.e. proliferation of precursor cells, migration and positioning of neuroblasts as well as neuronal maturation. Importantly, the development of distinct subtypes of cortical GABAergic interneurons was discretely impaired as a result of maternal inflammation. This translated into a reduction in cell numbers, redistribution across cortical regions and layers, changes in morphology and cellular properties. Furthermore, selective vulnerability of GABAergic interneuron subtypes was associated with the stage of brain development. Thus, we propose that maternally-derived insults have developmental stage-dependent effects which contribute to the complex etiology of cognitive impairment in the affected offspring.

## Introduction

Development of the fetal brain is highly influenced by the maternal environment. A multitude of factors such as maternal nutrition, stress, hormonal imbalance as well as the maternal immune status play key roles in shaping normal brain development^1^. Maternal inflammation is known to increase the risk for severe psychiatric disorders including schizophrenia, bipolar disorder, intellectual disability, anxiety, autism spectrum disorders (ASDs) and cerebral palsy^2,3^ with high societal costs^4^. Although the exact mechanism of adverse neurodevelopment upon maternal inflammation is not known, data from animal experiments and clinical observations suggest that the cytokine-related pro-inflammatory response in the mother contributes to disordered development of the fetal brain and predisposes the offspring to additional stressors during postnatal maturation of the brain^5–7^.

Cortical GABAergic interneurons arise during embryogenesis in the ganglionic eminences (GEs). Two of the three classes of GABAergic interneurons are generated in the medial ganglionic eminence (MGE), i.e., the parvalbumin (PV)- and somatostatin (SST)-expressing interneuron types, whereas the third class is generated in the caudal ganglionic eminence (CGE) and gives rise to a highly heterogeneous group of neurons that express serotonin receptor 3A (5HT3AR) and consists of vasoactive intestinal polypeptide (VIP), neuropeptide Y (NPY), reelin-positive and a few other subtypes^8–11^. In mice, cortical GABAergic interneurons are produced during middle-late gestation (E9.5-E18.5)^12,13^, and migrate tangentially from the GEs to the cortical plate, where they start migrating radially into prospective cortical layers^14,15^. Subsequently, a significant proportion of interneurons undergoes programmed cell death^16,17^, while the surviving cortical interneurons mature over two to three months in mice^9^.

Excitation-inhibition imbalance in the brain has been proposed as a major factor underlying the behavioral outcomes and cognitive decline associated with various neurodevelopmental disorders, including those associated with maternal inflammation^18–21^. Indeed, several studies show changes in activity and distribution of GABAergic interneurons in the cortex and hippocampus of the adult mouse brain upon maternal inflammation^22–24^. Such abnormalities are thought to occur due to perturbed brain development during early fetal or juvenile periods. Importantly, the development of neuronal circuits in the brain continues until late adolescence, up to 60-70 days postnatally in rodents and 20-25 years in humans^25^. Despite being acute, maternal inflammation can have long-lasting effects on several critical developmental processes such as precursor cell proliferation, neuronal migration and differentiation during embryogenesis, as well as postnatal neuronal maturation including neuronal survival, dendritic, synaptic and axonal pruning and synaptogenesis. Some studies have addressed the developmental impairment of principal cortical neurons upon maternal inflammation^6,26,27^, but surprisingly little is known regarding how development and maturation of cortical GABAergic interneurons are affected, and more importantly how those defects of GABAergic interneurons that are observed in adult animals^23,24,28^ arise during brain development. Abnormal development of cortical inhibitory circuits will have a significant impact on animal behavior leading to phenotypes resembling human psychiatric disorders as has been shown for a number of genetic mouse models^29–31^. Furthermore, as cortical GABAergic interneurons represent a diverse class of neurons with more than twenty subtypes^32^, various subtypes of GABAergic interneurons might be differentially affected by maternal inflammation contributing to the complex behavioral abnormalities of the offspring.

To study the impairment of cortical interneuron development due to maternal inflammation, we utilized a mouse model of maternal inflammation that involved injecting polyriboinosinic–polyribocytidilic acid (poly I:C), a synthetic polynucleotide that mimics viral double stranded RNA, at embryonic day 9.5 (E9.5). This early time point is equivalent to the first-trimester period in humans and represents a period of high vulnerability to neurodevelopmental disorders caused by maternal infections^33^. It also marks the onset of cortical GABAergic interneuron production from the MGE in mice^34^ as well as the pioneer Cajal-Retzius cells^35^. Poly I:C injection is a well-established model of maternal inflammation triggering an acute inflammatory response by activating pro-inflammatory cytokines including IL-2, IL-6, IL-8, IL-1β, IL-17a and TNFα^6,36^. It seems that these cytokines do not activate the embryonic microglia suggesting that maternal inflammation directly perturbs neuronal development^37,38^. Offspring born to poly I:C injected mice show marked behavioural deficits such as decreased pre-pulse inhibition, altered social interaction and cognitive decline^39,40^. Thus, to understand the effect of maternal inflammation on interneuron development, we undertook a detailed analysis of embryos and pups exposed to maternal inflammation. We found poly I:C to have an acute effect on GABAergic interneuron precursor proliferation, migration, positioning and maturation of GAD+ neuroblasts. The effect of maternal inflammation was interneuron subtype-specific and demonstrated differential vulnerability of interneuron subtypes to mother-derived insults.

## Materials and Methods

### Animal breeding and genotyping

All animal experiments were conducted in accordance with the guidelines of the National Animal Ethic Committee of Denmark. C57BL/6J (Janvier Labs), PV^Cre^ (017320, Jackson Labs), Ai9-tdTomato (007905, Jackson Labs) and GAD67-EGFP (Gad1^tm1.1Tama^)^41^ mice were used in this study. The mice were kept in IVC cages and housed under a reversed light cycle with food and water *ad libitum*. Heterozygous GAD67-EGFP mice were bred with wild-type mice. Females were kept with males until the vaginal plug was detected and then males were separated. Date of the vaginal plug was treated as E0.5, and embryos and pups were timed accordingly. Pups were weaned at P21 and littermates of the same sex were kept separately at 3-6 animals per cage.

EGFP+ animals were identified by PCR using the Accustart II Polymerase (QuantaBio). PCR conditions used were: 94°C 2min, followed by 35 cycles of 94°C 15s, 62°C 30s, 72°C 30s and final extension at 72°C for 5 min. The following primers (in 5’-3’ orientation) amplified a 345 bp region.

EGFP_F CCTACGGCGTGCAGTGCTTCAGC
EGFP_R CGGCGAGCTGCACGCTGCGTCCTC

### Induction of Maternal Immune Activation using poly I:C

Commercially prepared poly I:C (Sigma, P9582) was purchased and dissolved in phosphate buffered-saline (PBS) to give a 1 mg/ml stock solution. Two lots of poly I:C were used during the course of this study and no lot-specific effects of poly I:C were observed (Suppl. Figure 8, Suppl. Table 1).

Pregnant females at E9.5, E12.5 and E16.5 received a single tail intravenous (i.v.) injection of poly I:C equivalent to 5mg/kg body weight under mild restraint. Control females received an equal volume of PBS (no randomization regarding assignment to PBS or poly I:C injection). Information regarding rates of abortion and size of litters for poly I:C and PBS injected mice, as well as number of litters, animals and females/males used for each experiment is shown in Suppl. Table 2 and 3. Lack of litter-specific effects for PBS- or poly I:C injections are shown in Suppl. Figure 9 and Suppl. Table 4.

For experiments involving BrdU labelling of progenitors, 50 mg/kg of BrdU (Sigma, B5002) was chosen as in our previous studies^42,43^ and was injected intra-peritoneally (i.p.) into dams at E10.5, E14.5 and E17.5. Dams were sacrificed 2 hours after labelling and embryo were dissected and their heads collected in 4% paraformaldehyde solution.

### Plasma IL-6 estimation

Three hours-post poly I:C or PBS injection, blood samples were taken from the tail vein in EDTA pre-coated Eppendorf tubes. Following centrifugation (3000 rpm, 10 mins), the supernatant was collected in a separate tube and stored at −80°C when all samples were analyzed together. A Mouse IL-6 DuoSet ELISA kit (R&D Systems) was used to measure IL-6 levels. Absorbance was measured at 450 and 560 nm using a Glomax microplate reader (Promega). For analysis, readings at 560 nm were subtracted from those at 450 nm and normalized to blank controls. Values were obtained by linear regression of a plot of concentrations of known standards versus normalized absorbance values.

### Mouse behavior

PBS- and poly I:C-treated offspring (P35, male and female) were first acclimatized to the room and subsequently placed in the center of a circular open field (OF) of 1 m diameter. Movement was recorded for 10 mins using the Noldus Ethovision XT video tracking system v5 (Noldus Information Technology). Time spent in the central zone (0.7 m inner diameter) and periphery was extracted as were basic locomotor parameters such as total distance travelled, in total and in 1-min intervals and mean velocity.

Social interaction was tested immediately after the Open Field test. For this, two rectangular wire containers (6.2 cm × 7.8 cm × 12.4 cm, HxWxL) large enough to hold a mouse and allowing for social interaction were placed equidistant from the center of the open field. An unknown mouse was alternated between the two containers with the other kept empty. A region of 10 cm around the containers was treated as the social interaction zone, the remaining part of the arena as a non-social zone. Time spent by control and poly I:C mice in the social and non-social zones was extracted via Ethovision.

Acoustic startle reaction (ASR) and pre-pulse inhibition (PPI) were tested at 5 weeks of age as described^44^ in two chambers (San Diego Instruments, San Diego, USA) with 70 dB(A) white background noise. A piezoelectric accelerometer transduced displacement of mouse test tubes (Ø3.6 cm) in response to movements of the animal. Animals were acclimatized for 5 min in the tube before sessions started and ended with 5 startle trials of 40 ms 120 dB(A) bursts of white noise. In between, 35 trials were delivered in semi-randomized order (10 trials of 120 dB(A); 5 each of 4 pre-pulse + startle trials (pre-pulses of 72, 74, 78, and 86 dB(A)); 5 trials with only background noise). Tube movements were averaged over 100 ms following onset of the startle stimulus (AVG). The five AVGs for each pre-pulse intensity were averaged and used to calculate PPI, which was expressed as percent reduction in averaged the pre-pulse AVGs compared to the average of the 10 middle startle trials: %PPI = [1−(Pre-pulse + pulse/Pulse)] × 100.

### Perfusion, Sectioning and Immunohistochemistry

All steps were performed at ambient temperature unless otherwise noted. A description of antibodies used in this study is given in Suppl. Table 6.

Postnatal mice were anaesthetized with a combination of xylazine and ketamine injected i.p. This was followed by transcardial perfusion initially with cold PBS to flush out blood and then with cold 4% paraformaldehyde (PFA). Brains were then removed and post-fixed in 4% PFA overnight at 4°C before being stored in PBS with 0.01% sodium azide.

Brains were sectioned using a vibrating microtome (Leica, VT1000S) at a thickness of 50 μm and stored in PBS with 0.01% sodium azide at 4°C. Embryonic day (E) 10.5 to 14.5 embryos were fixed in 4% PFA overnight, dehydrated in 30% sucrose and sectioned at 25 μm thickness using a cryostat (Leica CM3050).

For staining, sections were blocked and permeabilized with 3% BSA (in 0.2% Triton X-100 containing PBS). Subsequently, they were incubated overnight at 4°C with appropriate primary antibodies. The following day, sections were washed and incubated with Alexa Fluor conjugated secondary antibodies for 2 h at room temperature. Nuclei were counterstained with DAPI (Sigma) and coverslips were mounted on slides with FluorSave (Merck).

BrdU labeling was done similar as before^45^ by treating with 1M HCl at 37 C for 30 min, followed by neutralization with 10mM Tris-HCl (pH8.5), and antibody labelling was carried out as described above. For BrdU and Nk×2.1/COUP-TFII double labelling, antigen retrieval was performed before BrdU labeling using sodium citrate solution (10mM) at 85°C for 15 min.

### Image acquisition and analysis

Images were acquired using a confocal microscope (Leica SP8, Leica Microsystems) and analyzed using ImageJ (NIH) and Imaris (Bitplane AG). After correcting for brightness and contrast, figures were prepared using Adobe Illustrator (Adobe Inc). Graph preparation and statistical analysis were carried out using Prism 7.0 (GraphPad).

Distribution of GABAergic interneurons across the cortex was analyzed in the region of interest (ROI) spanning from the upper edge of the corpus callosum or the subventricular zone to the pia at P1-60 old mice or E14.5-E17.5 old mice, respectively, and having 200-400 μm in width (medial-lateral) depending on age. The whole length of ROI was subdivided in 10 equal bins for E14.5-P9 ages, and in 6 layers at P15-60 when cortical layering could be distinguished based on DAPI staining. Presumptive motor and somatosensory regions in the developing brain are referred to as motor and somatosensory cortex, respectively.

The direction of migration at E17.5 was determined in the intermediate zone and cortical plate using ImageJ according to the direction of the major neurite expanding from the cell body of the neuroblast, and the angle of migration was measured between the vertical axis (from the ventricle to the pia) and the axis of the major neurite.

### Electrophysiology

Acute brain slices were prepared from P53-57 old mice. To determine an appropriate n-number, we calculated before performing the experiments the required sample size to reach significance (p < 0.05) if the observed effect was more than 20%. The power analysis was based on means and distributions of data from our previous electrophysiological experiments. Using G*Power for this analysis (α-error = 0.05; Power (1−β error) = 0.85), we determined that n-number of 17 per group as required. We recorded from a slightly higher number of cells (26 cells from 3 PBS-injected mice and 25 cells from 4 poly I:C-injected mice), and 2-3 neurons were recorded per brain slice. Only basket cells were recorded and few chandelier cells were excluded from the analysis. To this end, mice were deeply anesthetized with Isoflurane (3%) and the brains quickly removed and dissected in ice-cold sucrose-containing ACSF (212 mM sucrose, 0.02 mM CaCl_2_, 7mM MgCl_2_, 3 mM KCl, 1.25mM NaH_2_PO_4_, 26mM NaHCO_3_, 10mM d-glucose oxygenated with carbogen). 250μm thick coronal brain slices were cut with the help of a tissue slicer (Leica VT1200S, Wetzlar, Germany) in ACSF (25 mM d-glucose, 125 mM NaCL, 1.25mM NaH_2_PO_4_, 26mM NaHCO_3_, 2.5 m M KCl, 2 mM CaCl_2_, 1mM MgCl_2_ oxygenated with carbogen).

Brain slices were visualized under an upright Olympus BX51WI microscope fitted with a 40x water-immersion objective (LUMPlan FI/IR, NA 0.8w; Olympus, Japan) with infrared optics. Whole-cell recordings were obtained from single neurons in layer 2/3 of the somatosensory cortex under visual guidance. On average, 2-3 neurons were recorded per brain slice. Borosilicate glass pipettes (3-5MΩ) were pulled with a micropipette puller (Sutter Instruments, Novato, CA, USA) and filled with intracellular solution (130 mM K-gluconate, 10 mM HEPES, 10 mM phosphocreatinine- Na, 10 mM Na-gluconate, 4 mM ATP-Mg, 0.3 mM GTP, 4 mM NaCl; pH 7.2). Intrinsic membrane properties and firing patterns were recorded in current clamp mode with 50 pA current steps ranging from −50 pA to 900 pA. Intrinsic membrane properties and firing properties were analyzed with IGOR Pro (WaveMetrix, USA) and Microsoft Excel (Microsoft, USA).

### Western blot

Mice were euthanized by cervical dislocation and brains were quickly removed. The cortex was dissected in PBS on ice and stored at −80°C until use. Cortices were homogenized using a Dounce tissue grinder set (Loose followed by Tight pestle, Sigma) in ice cold 300 μl RIPA buffer supplemented with Complete EDTA-free Protease inhibitor (Roche). Lysates were incubated for 10 min at 4°C and collected by centrifugation at 5000 × *g* for 10 min. Protein concentration was determined using the bicinchoninic acid (BCA) kit (Pierce) and BSA as a standard. Equal amount of SDS sample buffer, Laemmli 2x concentrate (Sigma) was added to each lysate and boiled for 5 min. Samples were separated by 15% SDS-PAGE gel and transferred onto the 0.45 μm pore size Immobilone-FL Membrane (Millipore) for 1 h at 20 V. Membranes were blocked in 3% BSA (Applichem) in TBS-T (TBS plus 0.1% Tween 20) for 1 h at room temperature and then incubated overnight at 4°C with primary antibody: mouse anti-GAPDH (1:5000, Abcam) or rabbit anti-parvalbumin (1:10000, Swant) diluted in blocking solution. After washing with TBS-T, membranes were incubated with secondary horse anti-mouse HRP-conjugated (1:20,000; Vector laboratories) or anti-rabbit HRP-conjugated (1:20,000; GE Healthcare) in blocking solution for 1 h at room temperature. Detection was performed using the ECL chemiluminescence reagent (Pierce) and X-ray films (AGFA).

### Statistical analysis

Statistical analysis of the data was performed with Prism (GraphPad software, USA). Normality of data distribution was analyzed by D’Agostino and Shapiro-Wilk’s tests. Normally distributed data was analysed using the Student’s t-test (for 2 groups) or by 1-way ANOVA test (for more than 2 groups). Welch’s correction was applied when standard deviations of the two groups differed from each other. For simultaneous comparison of 2 parameters between 2 and more groups, we used 2-way-ANOVA and the Bonferroni post-hoc test. Equality of variances was analyzed using Dunnett’s test. A linear mixed model (LMM) analysis was used to analyse electrophysiology results in Figure 5b,d,f. We extended the LMM analysis to address the issue of a shared intrauterine environment in multiparous species^46^, especially applying it to outcomes of the histological analysis (Suppl. Table 5). For multiple comparisons, p-values from LMM analysis were corrected with Benjamini-Hochberg method.

To determine an appropriate number of animals, we estimated sample size based on our previous experiments and on similar experiments published by other groups. The investigators were not blinded to allocation during experiments and outcome assessment. Females and males were analyzed together, and all the graphs represent joint sex data unless otherwise stated. For all behavior experiments, sex-related differences were analyzed by 2-way ANOVA with exposure and sex as factors.

Basal startle and habituation to the first five 120 dB noise bursts and PPI were analyzed by 2-way ANOVA with poly I:C exposure and sex as factors; for basal startle habituation with the first five 120 trials as the repeated measure, for PPI with prepulse level (72, 74, 78, 86 dB(A)) as the repeated measure.

## Results

### Maternal inflammation model in transgenic mice that labels all cortical GABAergic interneurons

In order to reveal how the development of cortical GABAergic interneurons is affected by maternal inflammation, we utilized the GAD67-EGFP knock-in transgenic mouse line that labels all GABAergic interneurons in the cortex by EGFP^41^. Maternal inflammation was induced by injecting poly I:C in pregnant dams at E9.5 corresponding to the early first trimester of pregnancy in humans^47^; a period of highest risk of induction of neurodevelopmental disorders caused by maternal infections^33^. This period also marks the onset of cortical GABAergic interneuron production in mice which starts in the MGE at E9.5^34^. We confirmed the induction of inflammation by an increase in maternal plasma IL-6 levels three hours after injection of poly I:C both in wild-type and EGFP+ females (Figure 1a). To validate that poly I:C mouse model in our hands has a similar functional impact on the offspring as previously described^40,48^, we performed behavioral experiments on control and poly I:C-affected offspring testing for social interaction and memory, open field and startle (including pre-pulse inhibition (PPI)) behavior. Control mice on average spent significantly more time interacting with the unfamiliar mouse in comparison to the inanimate object (the empty container), while offspring affected by maternal inflammation did not have such a preference (Figure 1b). Control offspring also interacted with the unfamiliar mouse for longer durations than those affected by maternal inflammation (Figure 1c). In addition, offspring affected by maternal inflammation displayed signs of hyperactivity observed as an increase in the total distance travelled and in running speed in the open field test (Figure 1d,e). For the basal startle reaction and the habituation to the first five 120 dB noise bursts, there were no statistically significant differences owing to poly I:C-exposure; the startle reaction in males was significantly stronger than in females (data not shown). ANOVA analysis of PPI, with poly I:C exposure and sex as factors and level of prepulse intensity as the repeated measure (72, 74, 78 and 86 dB(A)), showed significant interaction between poly I:C exposure and prepulse level (p=0.049), whereas interaction between prepulse level and sex had p>0.05 (there was also no three-way interaction between prepulse level, poly I:C exposure and sex). Separate analysis of PPI at each prepulse level indicated that the statistically significant interaction was due to differences at the 72 dB prepulse level (Suppl. Figure 1a), where exposed offspring showed reduced inhibition compared to control offspring (Figure 1f). Overall, we did not observe marked differences in behavior between males and females (except for total distance travelled in the open-field test) (Suppl. Figure 1b-j). Thus, maternal inflammation triggered by poly I:C injection led to impaired social behavior, increased anxiety and sensorimotor gating deficits in wild-type mice. These phenotypes were similar to those observed in previous studies of poly I:C injected mice^5,6,22,49–53^.

**Figure 1:**
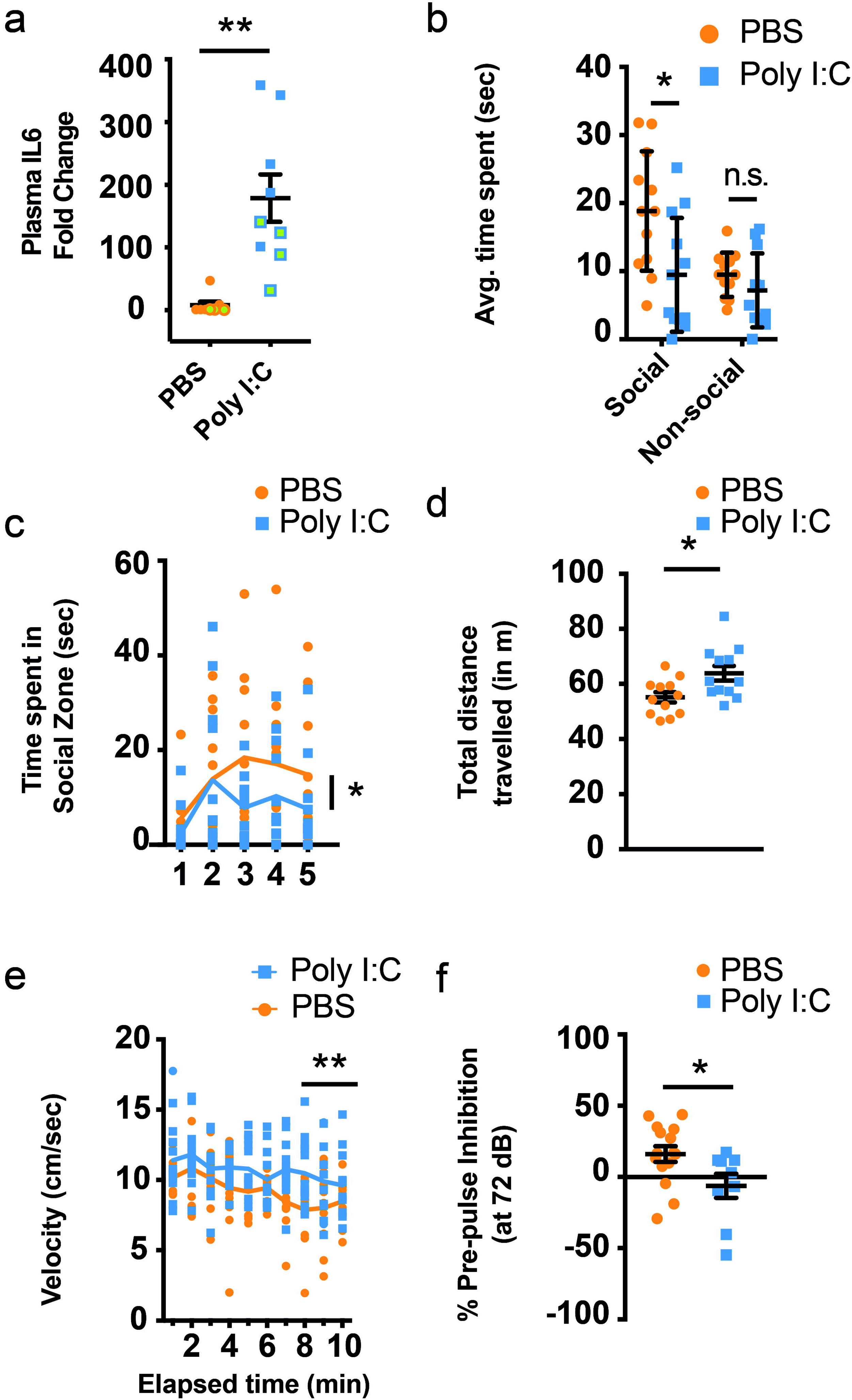
Poly I:C induces rapid upregulation of IL-6 in maternal plasma and behavioral impairments in maternal inflammation affected offspring. (a) Upregulation of IL-6 protein in maternal plasma 3 hours after i.v. injection of poly I:C (n=8, 9 for PBS and poly I:C respectively, 5 wild-type and 3,4 GAD67-EGFP mice respectively, labeled by filled and green circles/squares, respectively). (b,c) Prenatal exposure to maternal inflammation results in impaired social-zone preference (b) (n=12 each), as well as a decrease in time spent in social zone (c) (F=3.23). (d,e) Mice exposed to maternal inflammation during gestation present signs of hyperactivity as seen by total distance travelled (d) (n=12 each) and speed of running over a 10 minute period (n=12 each). (f) Maternal inflammation during early gestation induces schizotypal effects in offspring that can be gauged by a decreased pre-pulse inhibition (at 72dB) (n=15 and 9, for PBS and poly I:C, respectively). Comparison of means by Student’s t-test in (a,d,f), t-test with Holm-Sidak correction for multiple comparison in (b) and 2-way ANOVA in (c,e) (mean±SEM are shown, p-values denoted as asterisks with <0.05 shown as * and <0.005 shown as **).

### Maternal inflammation impacts early stages of cortical GABAergic interneuron development

While the effect of maternal inflammation on offspring behavior has been well documented, there is little information about how maternal inflammation affects the development of inhibitory circuits in the cortex that might underlie abnormal animal behavior. In the subsequent analyses, we focused mainly on the developing somatosensory and motor cortices, since these are the best characterized cortical areas in mice that are evolutionary conserved in humans and individuals with schizophrenia frequently show anatomical and functional impairments in motor and sensory areas^54–56^. We therefore investigated the effect of poly I:C (injected at E9.5) on the early stages of interneuron development. Cortical GABAergic interneurons originate between E9.5 and E18.5 in the GEs^12,13^ from where they migrate tangentially into the developing cortex. They enter the cortical plate at two principal sites – a superficial stream along the marginal zone just below the pia, and a deeper stream at the intermediate zone/ sub-ventricular zone (IZ/SVZ)^57,58^. Once in the developing cortical plate, GABAergic interneurons switch to a radial mode of migration and invade the cortical plate (Figure 2a). Strikingly, we observed a clear effect of maternal inflammation on cortical GABAergic interneurons early during their development, since already at E14.5 fewer EGFP+ neuroblasts were observed in the IZ of poly I:C exposed embryos at the regions of the developing cortex corresponding to the motor and somatosensory cortices (Figure 2b,c). In maternal inflammation affected embryos, there were no evidence in disturbance of stream of neuroblasts migrating from the GE to the cortical plate, and we did not find large assembles of neuroblasts that were stuck during migration (Suppl. Figure 2a). Additionally, the density of EGFP+ neuroblasts in the SVZ did not change in response to maternal inflammation (Suppl. Figure 2e). To study differences in migratory patterns of GABAergic interneurons, we chose E17.5 time-point when the majority of interneurons reach the developing cortex and start to migrate radially within the cortex. We divided the cortex into 10 equally sized bins (see Materials and Methods) and quantified the number of EGFP+ neuroblasts in each. The thickness of the cortical plate, IZ and MZ did not show any changes in E14.5 or E17.5 embryos due to exposure to maternal inflammation (Suppl. Figure 2b,c). Remarkably, maternal inflammation resulted in large-scale redistribution of GABAergic interneurons with an evident reduction in density of EGFP+ neuroblasts in the regions corresponding to the motor and somatosensory cortices of the developing cortex at E17.5 (Figure 2d,d’). Quantification of densities per bin showed that pups exposed to maternal inflammation had fewer EGFP+ neuroblasts in the superficial layers of the cortical plate (Figure 2e, bins 3-4 for M1 and bin 4 for S1), which also resulted in a decrease in total neuroblast density across the cortical plate and intermediate zone (Figure 2f). However, while the developing somatosensory cortex showed an additional increase in percentage of EGFP+ neuroblasts to stack in the middle layers, a similar effect was not seen in the motor cortex (Figure 2d,d’,e, bin 5), indicating a differential effect of maternal inflammation on different cortical areas.

**Figure 2:**
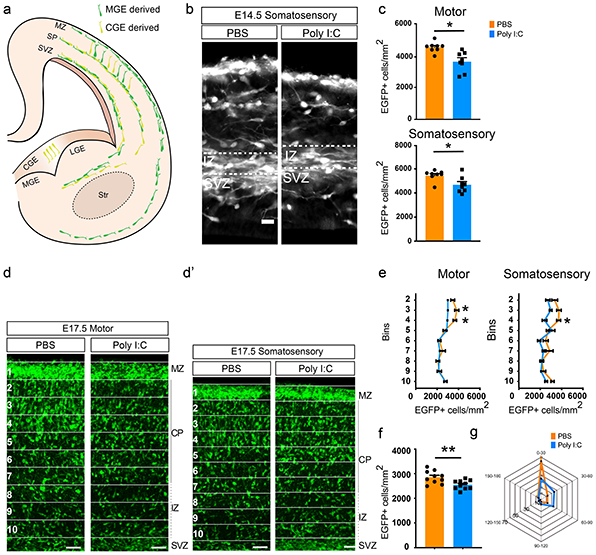
Early migratory deficits of GAD-EGFP+ neuroblasts due to maternal inflammation. (a) A schematic overview of the migratory routes taken by GAD+ neuroblasts during embryonic development^57,58^. (b,c) Reduction in the density of EGFP+ neuroblasts migrating into the cortical intermediate zone at E14.5 as a result of maternal inflammation. Panel (b) shows somatosensory region of the developing cortex. Dotted box denotes the region of counting. Quantifications from the dotted boxes are shown in (c) (motor cortex: n=8 each; somatosensory cortex: n=8 each). (d,e) Superficial regions of the cortical column show a reduction in density of EGFP+ neuroblasts due to maternal inflammation. Cortical columns from developing motor (d) and somatosensory (d’) cortical regions were divided into 10 equal-sized bins and the number of EGFP+ cells counted in each (with the exception of bin 1). Bin 10 represents the upper margin of the IZ. Densities in the graphs (e) show a reduction in bins 3-4. (f) Embryos exposed to maternal inflammation showed a decreased total density of migratory neuroblasts at E17.5. EGFP+ cells were counted in the cortical plate and intermediate zone across bins 2-10 (n=10 each). (g) Radar plot showing that a greater percentage of neuroblasts in the cortical plate migrate laterally due to maternal inflammation as assessed by the angle of leading process. Angles were grouped into 30° bins and relative percentages were plotted. Each spoke represents 10%. Comparison of means by Student’s t-test in (c,f), t-test with Holm-Sidak correction for multiple comparison in (e) (mean±SEM are shown, p-values denoted as asterisks with <0.05 shown as * and <0.005 shown as **). Scale bars: 50 μm

To account for the effect on distribution of GAD+ neuroblasts caused by acute maternal inflammation we analyzed the directionality of neuroblast migration in the IZ and cortical plate at E17.5. While GAD+ neuroblasts in the cortex of control offspring predominantly migrated towards the pia, their direction in inflammation-affected cortices was significantly different with a greater fraction migrating laterally (Figure 2g and Suppl. Figure 2d).

### Maternal inflammation impairs positioning of GABAergic interneurons in the developing cortex

As interneuron migration continues until the second postnatal week, we chose to analyze whether these initially observed differences continue to affect the positioning of interneurons in the postnatal cortex. We hence studied several postnatal stages from P3 to P30 to understand how maternal inflammation can lead to a disordered arrangement of GABAergic interneurons in the mature cortex.

At P3 and P6, the difference in GAD+ cell density had amplified to cover the entire length of the cortical plate with the exception of the lowermost bins (bins 9-10, Figure 3a,b, Suppl. Figure 3a,b). Furthermore, the magnitude of the difference was consistently greater in the prospective somatosensory cortex than the motor cortex. Interestingly, the relative distribution of GAD+ neuroblasts across bins was similar in pups exposed to PBS or poly I:C (Suppl. Figure 3d,e). This suggests that the decrease in GAD+ neuroblasts is equally distributed across the whole length of the cortex. By P9, while the reduction in density persisted in the somatosensory cortex, much lower difference could be observed in the motor cortex (Figure 3c, Suppl. Figure 3c) indicating a differential effect of maternal inflammation on neuroblasts destined for distinct cortical regions.

**Figure 3:**
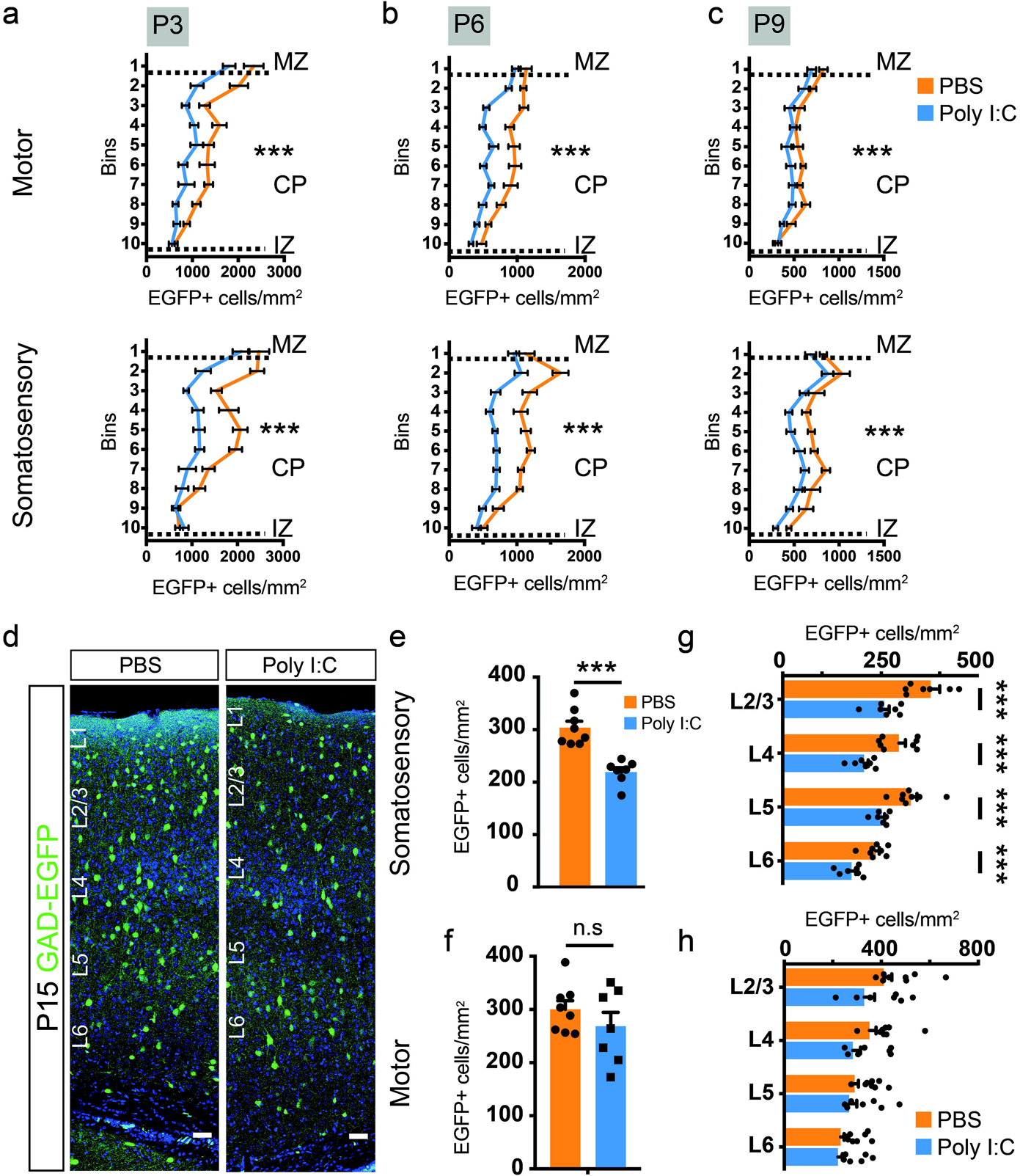
Maternal inflammation causes large-scale impairment in the distribution of GAD-EGFP+ neuroblasts and mature interneurons in the cortex. (a-c) Graphs showing a decrease in density of EGFP+ cells measured by counting EGFP+ cells in 10 equally sized bins in the cortical plate of the presumptive motor and somatosensory cortices. Dotted lines represent the margins of IZ, CP and MZ respectively (P3 motor: F=21.85, P3 somatosensory: F=24.1; P6 motor: F=21.72, P6 somatosensory: F=19.59; P9 motor: F=12.22, P9 somatosensory: F=13.77) (d-h) Significant reduction in the density of interneurons in the somatosensory but not in the motor cortex at P15. EGFP+ cells were quantified in the total cortical thickness as well as in each cortical layer. Panels in (d) show representative images of regions used for quantification. Graphs show the total (e,f) (n=8 and 7, for PBS and poly I:C, respectively) as well as the layer-wise (g,h) reduction in EGFP+ cell density. Comparison of means by 2-way ANOVA in (a-c), Student’s t-test in (e,f) and t-test with Holm-Sidak correction for multiple comparison in (g,h) (mean±SEM are shown, p-values denoted as asterisks with <0.0005 shown as ***). Scale bars: 50 μm (d).

Cortical GABAergic interneurons are overproduced during embryogenesis, and approximately 30-40% of them subsequently undergo apoptosis within the second postnatal week^16^. We therefore investigated whether an increase in programmed cell death of cortical interneurons due to maternal inflammation could explain in part the reduction of their number in the cortex of maternal inflammation-affected mice. To this end, we counted activated caspase-3 and EGFP-expressing cells in several sections covering the motor and somatosensory cortices. At E17.5, P3 and P6, we observed a similar number of activated caspase-3+ (Casp3+) as well as EGFP+ Casp3+ cells in control pups and pups exposed to maternal inflammation (Suppl. Figure 4a,b). Thus, activation of the maternal immune system during early gestation did not augment apoptosis in the developing cortices of pups exposed to maternal inflammation.

To follow up the effect of maternal inflammation during postnatal brain maturation, we further studied the density and positioning of interneurons at P15. Similar to P9, the density of GAD+ interneurons in S1 was severely reduced (Figure 3d,e), whereas no significant changes were observed for M1 (Figure 3f). Analysis of the layer-wise distribution of interneurons showed an equal effect of maternal inflammation across the whole cortical length in the somatosensory cortex (Figure 3g), but lack of reduction in GABAergic interneuron density in the motor cortex (Figure 3h). We hence show that an acute induction of inflammation in pregnant dams impairs migration and final positioning of GAD+ neuroblasts with the effect being more pronounced in the somatosensory than in the motor cortex.

### Differential effect of maternal inflammation on distinct subtypes of cortical GABAergic interneurons

While functional impairment of cortical GABAergic interneurons has been proposed to be one of the major factor contributing to imbalance between excitation and inhibition in patients with schizophrenia and other psychiatric disorders, most of the attention has been directed towards dysfunction of PV+ GABAergic interneurons^24,59,60^. However, cortical GABAergic interneurons are highly heterogeneous, and PV+ interneurons represent only one class of interneurons^8^. Therefore, the differential impact of maternal immune activation on development and maturation of various interneuron subtypes remains to be elucidated.

To this end, we studied the effect of maternal inflammation on the organization of the four largest interneuron populations, i.e. those that express PV, SST and VIP and CR (Figure 4a,b, Suppl. Figure 5a,b). In the somatosensory cortex, the previously observed decrease in the density of GAD+ interneurons stemmed from a decrease in both PV+ and SST+ interneurons (Figure 4a-d). Interestingly, there was a clear layer-specific effect of maternal inflammation in the distribution of PV+ and SST+ interneurons. Thus, the decrease in number of PV+ interneurons was restricted to layer 4 at P15 (Figure 4c), whereas the reduction of SST+ interneurons was more pronounced across all layers, albeit statistically significant only in layers 2-4 at P15 (Figure 4d). There was no effect of maternal inflammation on the distribution of CGE derived VIP+ and CR+/SST− interneurons at P15 (Suppl. Figure 5a,b). At P60, the densities of PV+ and SST+ interneurons were not statistically significant between maternal inflammation-affected and control animals even though an appreciable difference could be seen in the case of L4 SST+ interneurons (Figure 4e,f). This is likely caused by the non-uniform expansion of cortical areas during synaptogenesis and gliogenesis^61^. In the motor cortex, there was no significant effect on distribution of interneuron subtypes at P15 and P60 (data not shown). Similar numbers of PV or SST+ interneurons were also observed in the prefrontal cortical areas^62^, namely cingulate, prelimbic and secondary motor cortex (M2), of PBS and poly I:C exposed mice (Suppl Figure 5a,b).

**Figure 4:**
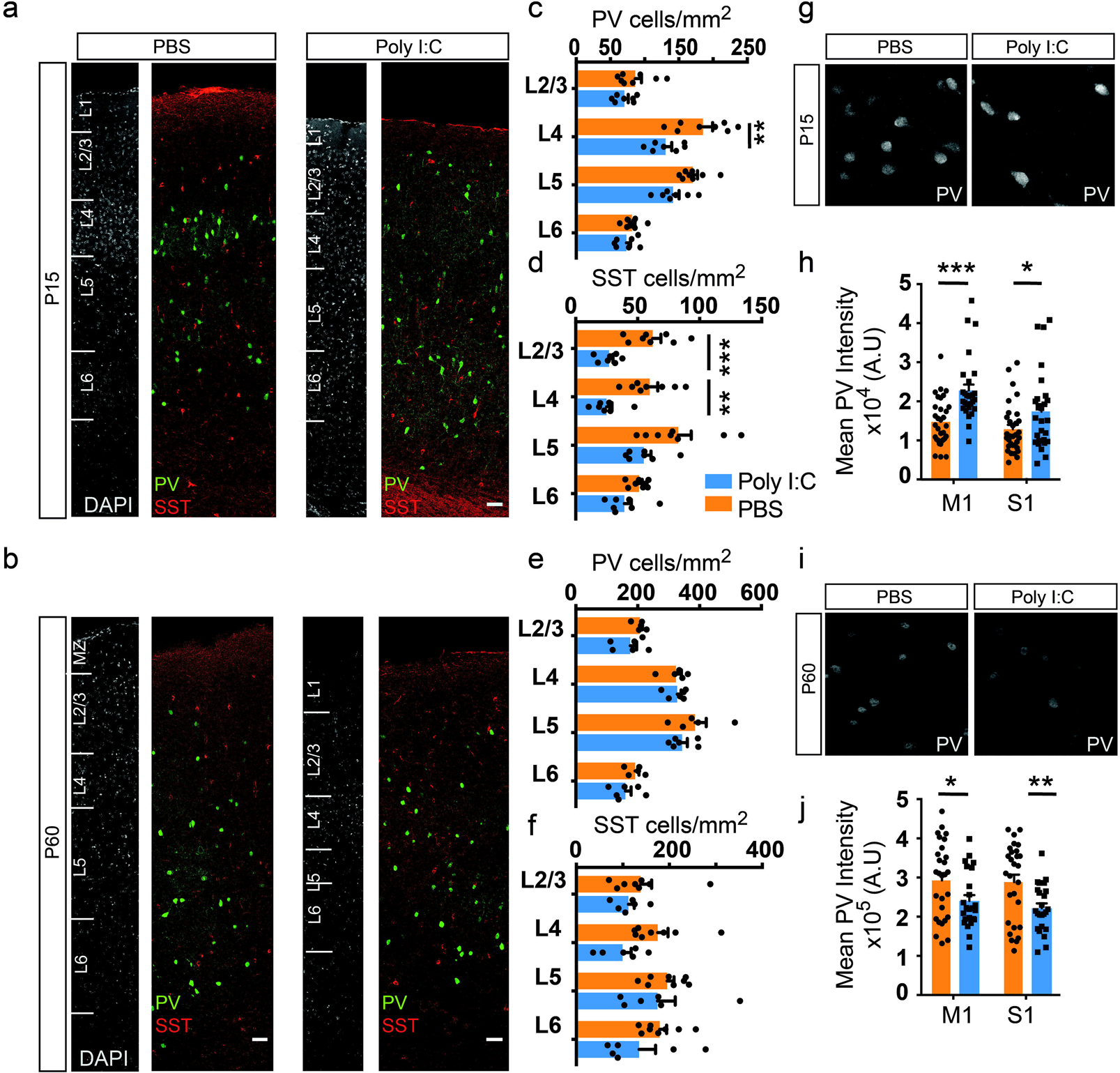
Interneuron subtype-specific effects of maternal inflammation. (a-f) Maternal inflammation resulted in a decreased density of PV and SST expressing interneurons at P15 but not at P60. Panels show representative images from P15 (a) and P60 (b). A significant decrease in PV+ interneurons could be seen in L4, while the difference was more pronounced in L2-4 for SST+ interneurons (c,d) (n=8 and 7, for PBS and poly I:C, respectively). An appreciable but not statistically significant decrease could be seen at P60 (e,f). (g-j) Reciprocal changes in PV expression between P15 and P60 due to maternal inflammation. At P15, PV+ interneurons in both motor and somatosensory cortical regions in maternal inflammation-exposed animals showed an increased expression of PV (h) (n=8 and 7, for PBS and poly I:C, respectively) measured by the signal intensity in immunolabeled images (g). By P60 (i), this effect had reversed and instead a decreased expression was observed in both motor and somatosensory cortical regions (j) (n=6 each). Statistical analysis by t-test with Holm-Sidak correction for multiple comparison in (c-f,h,j) (mean±SEM are shown, p-values denoted as asterisks with <0.05 shown as *, <0.005 shown as ** and <0.0005 shown as ***). Scale bars: 50 μm (a,b).

In addition to the reduced interneuron density, a reduction in PV expression in the cortex and hippocampus has been described in several studies for adult mice^21^. While we confirmed that maternal poly I:C exposure led to a similar reduction in PV expression in both M1 and S1 cortical regions at P60 (Figure 4i,j, Suppl. Figure 6), surprisingly, PV+ interneurons in both regions had an excess of PV expression at P15 (Figure 4g,h). As parvalbumin is an important calcium binding protein in neurons^63^, and its expression positively correlates with neuronal activity in the developing cortex^64^, the developmental impairment in calcium buffering capacity in PV-expressing neurons might contribute to their functional impairment. Previous studies have shown maternal immune activation to be detrimental to PV+ interneurons of the medial prefrontal cortex leading to decreased GABAergic transmission onto pyramidal neurons^22^. We thus assessed the morphology and functional properties of PV+ interneurons in the superficial layers of the somatosensory cortex in control and animals prenatally exposed to maternal inflammation at P53-57. PV+ interneurons were labelled by Cre-recombinase dependent tdTomato expression in Ai9 reporter mouse (*PV^Cre^; Ai9*) and were assessed for intrinsic electrophysiological properties using whole-cell patch-clamp recordings of fluorescent neurons in acute brain slices (Figure 5a-i). The maximum firing frequency of PV+ interneurons was reduced in poly I:C-exposed animals (Figure 5a,b) reflecting an overall slower spiking rate due to maternal immune inflammation. At smaller current injections, no difference could be seen (Figure 5c). Additionally, the resting membrane potential and input resistance in poly I:C-exposed animals was not found to be significantly different in comparison to controls (Figure 5d,e). To investigate if the changes in electrophysiological properties are accompanied by anatomical changes, we analyzed the morphology of biocytin-filled PV+ interneurons (Figure 5f). There was no significant effect of maternal inflammation on the length of dendritic filaments (Figure 5g) and branch depth (Figure 5h). However, Sholl analysis of the reconstructed interneurons showed a significant reduction in overall dendritic tree complexity of the maternal inflammation-affected PV+ interneurons (Figure 5i). These results indicate that acute maternal immune response affects morphology and electrophysiology of GABAergic interneurons.

**Figure 5:**
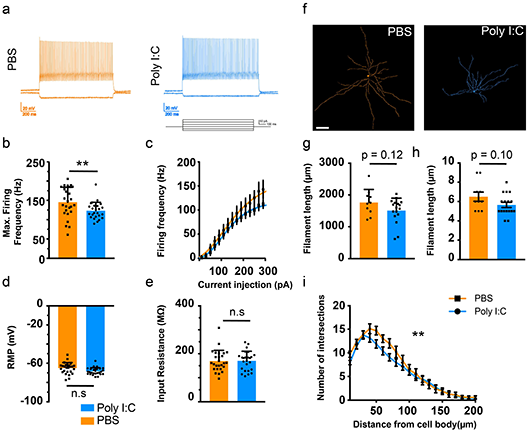
Impaired physiology and morphology of fast-spiking basket cells due to maternal inflammation. (a-e) Maternal inflammation resulted in altered intrinsic physiological properties of PV+ fast-spiking interneurons. Representative traces are shown in (a). Maternal inflammation affected cells showed a decrease in maximum firing frequency (b) (PBS: 145.1Hz±7.61, Poly I:C: 123.1Hz±4.36, n=25 cells each). (c) Changes in firing frequency were only evident at higher current injections and the resting membrane potential (RMP) and input resistance did not show any change (d,e). (f-i) Altered morphological complexity of PV+ fast-spiking interneurons was also observed by biocytin back-filling and reconstruction (f). Differences in filament length and branch depth (g,h) did not reach p-value <0.05, while Sholl analysis showed a decreased complexity (i) (F=179.7). Linear Mixed Model analysis in (b-e), comparison of means by Student’s t-test (g,h), and 1-way ANOVA in (i) (mean±SEM are shown, p-values denoted as asterisks with <0.05 shown as * and <0.005 shown as **). Scale bar: 30 μm (f).

### Timing of maternal inflammation affects discreet pools of interneuron progenitors

One of the most remarkable effects of maternal inflammation on development of the GABAergic interneurons was the decrease in the density of EGFP+ neuroblasts as early as E14.5 when maternal immune activation was induced at E9.5 (Figure 2b,c). As cortical GABAergic interneurons are generated by precursor cells mainly in the MGE and CGE between E9.5 and E18.5^12,13^, acute maternal inflammation might affect the generation of GABAergic interneurons. To investigate this, GAD67-EGFP pregnant mice were injected with poly I:C at E9.5, E12.5 and E16.5 and proliferating cells were labelled using a 2-hour pulse of the thymidine analog, 5-bromo-2’-deoxyuridine (BrdU), at E10.5, E14.5 and E17.5 (Figure 6a,e,h). By co-labeling interneuron precursors for BrdU along with Nkx2.1 (MGE-specific marker^34,65^) or COUP-TFII (CGE-specific marker^66^), we sought to ascertain differences in proliferation of interneuron precursors due to maternal inflammation.

**Figure 6:**
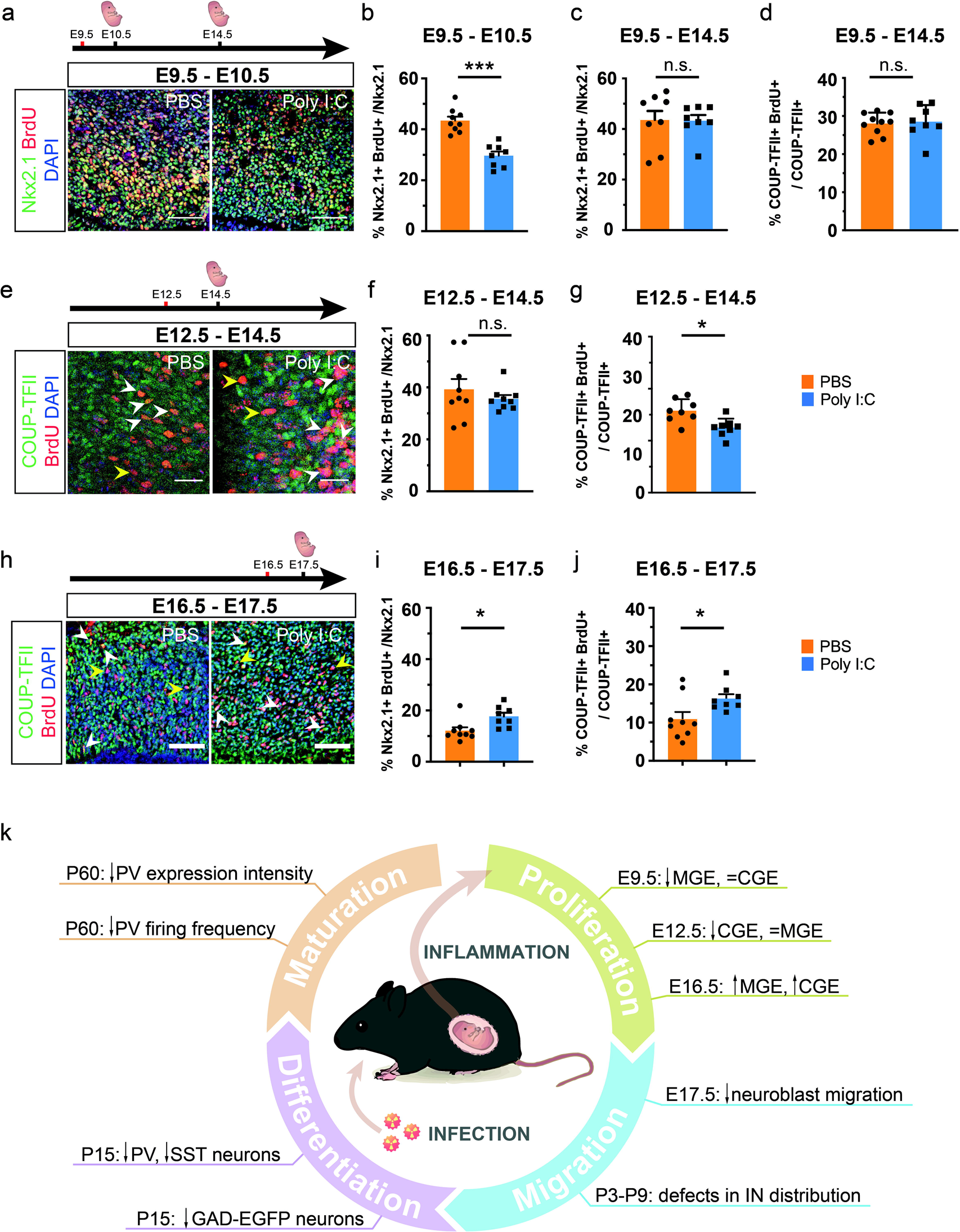
Acute and developmental stage-dependent impairment of progenitor proliferation due to maternal inflammation. (a-c) Maternal inflammation had acute effect on proliferation of MGE-derived Nkx2.1+ progenitors. Dams were injected with poly I:C at E9.5 and embryos assessed at E10.5 after a 2-hour BrdU pulse-chase (a-b). Quantification of proliferating Nkx2.1+ BrdU+ cells showed that maternal inflammation resulted in reduction in Nkx2.1+ proliferating cells at E10.5 (b) (n=9 and 8, for PBS and poly I:C, respectively) that did not sustain until E14.5 (c). COUP-TFII+ progenitors remained unaffected when studied at E14.5 (d). (e-g) Dams were injected with poly I:C at E12.5 and embryos assessed at E14.5 after a 2-hour BrdU pulse-chase. Proliferation of CGE-derived COUP-TFII+ progenitors was reduced after maternal inflammation at E12.5 (e,g) (n=9 and 8, for PBS and poly I:C, respectively) that correlated with the later neurogenic window of CGE-derived progenitors. In contrast, proliferation of Nkx2.1+ MGE-derived progenitors was unaffected at E14.5 (f). (h-j) Late maternal inflammation at E16.5 led to an increase in proliferation of both COUP-TFII+ (h,j) and Nkx2.1+ (i) progenitors (n=8-9 for each PBS and Poly I:C). (k) Schematic representation of the findings in this manuscript. Maternal inflammation affects interneuron development at multiple stages such as proliferation, migration, differentiation and maturation leading to increased vulnerability to mental disorders. Central artwork modified from^90^ Comparison of means by Student’s t-test in (b-d,f,g,I,j) (mean±SEM are shown, p-values denoted as asterisks with <0.05 shown as *, <0.005 shown as ** and <0.0005 shown as ***). Scale bars: 50 μm (a,e,h).

By injecting polyI:C at E9.5 and labelling progenitor proliferation at E10.5, we found a ~25% decrease in the percentage of Nkx2.1+ precursors that were co-labelled with BrdU in embryos exposed to maternal inflammation in comparison to non-exposed mice (Figure 6a-b), which correlates very well with a decrease in the number of MGE-derived PV+ and SST+ interneurons in the postnatal cortex. In contrast, both exposed and non-exposed mice showed equal rates of proliferating Nkx2.1+ progenitors five days post treatment, i.e. at E14.5 (Figure 6c). This indicates an acute effect of maternal inflammation on the cycling of interneuron progenitors. Furthermore, in keeping with the acute response of progenitors, maternal inflammation exhibited no effect on the proliferation of COUP-TFII+ progenitors over the same period (Figure 6d), which might be explained by the fact that the majority of CGE-derived interneurons are born after E12.5^12^. This correlated with a lack of an effect on CGE-derived VIP+ and CR+/SST− interneurons in the somatosensory cortex at P15 (Suppl. Figure 5d-f), thus indicating a subtype-specific effect of maternal inflammation on interneuron progenitors.

To investigate whether a later induction of maternal inflammation would affect both CGE-derived (born mainly between E12.5-18.5^12^) and MGE-derived interneurons, we injected dams with poly I:C at E12.5. As with the previous analysis, a 2-hour pulse of BrdU was given before collecting the embryonic brains for analysis at E14.5 (Figure 6e). Interestingly, we found no difference in the cycling Nkx2.1+ progenitor population at this stage between the two groups (Figure 6f). On the other hand, we observed a ~20% decrease in COUP-TFII+ progenitors that were co-labelled with BrdU (Figure 6g). Despite the decrease in proliferation during embryogenesis, there were no significant reduction in the number of CGE-derived VIP+ and CR+/SST− interneurons in the somatosensory cortex at P15, although we noticed that L5 VIP+ interneurons are virtually absent in maternal inflammation-affected offspring (Suppl. Figure 5g-i). Such minor effects can be explained by the relatively low production of VIP+ and CR+ subtypes at E12.5^13^. In line with our hypothesis of a developmental stage-dependent effect of poly I:C, we did not observe a change in the number of PV+ and SST+ interneurons due to poly I:C exposure at E12.5 (Suppl. Figure 5j,k). We also demonstrated that the effect of maternal inflammation was confined to the SVZ, since quantification of BrdU+ proliferating cells in the ventricular zones (VZ) of MGE and CGE after poly I:C injections at E9.5 and E12.5, respectively, did not show any significant differences from PBS injections (Suppl. Figure 7a,b).

Finally, as CGE interneuron production peaks at E16.5^12^, we injected poly I:C at this stage in order to capture the maximal effect of maternal inflammation on CGE-derived interneurons (Figure 6h). Surprisingly, and in contrast with the earlier time points, 24 hours later (i.e. at E17.5), we found a marked increase in dividing Nkx2.1+ MGE and COUP-TFII+ progenitors (~37% and ~70% respectively) (Figure 6i,j) indicating that maternal inflammation has an opposing effect at later stages of embryonic development. Therefore, our results show that progenitors of GABAergic interneurons exhibit differential vulnerability to maternal inflammation that depends on the stage of embryonic brain development.

## Discussion

Despite significant evidence showing that maternal immune activation during pregnancy leads to cognitive dysfunction in the offspring that might be triggered by excitation-inhibition imbalance, little is known about how the development of inhibitory system in the brain is affected. Here we show that maternal immune activation results in multiple “hits” on development of GABAergic neurons, thus affecting proliferation of precursors, migration and positioning of neuroblasts as well as their maturation (see summary in Figure 6k).

Genetic and environmental factors that affect cortical development provide important insights into the cellular and molecular basis of neurodevelopmental disorders. Available epidemiological and clinical findings^67^ point towards a link between maternal infections during gestation and increased risk of developing a mental disorder due to impaired cortical development. While the precise mechanism of action is elusive, an increase in maternal cytokine levels is thought to lead to the transmission of maternal inflammation to the fetal brain^3^. However, the developmental changes occurring upon maternal inflammation in the fetal brain and during postnatal brain maturation have been understudied owing to ethical and technical challenges in humans. Thus, rodent models of maternal inflammation provide an excellent alternative to study the effects of an acute inflammatory insult on brain development. Accordingly, poly I:C and lipopolysaccharide (LPS) have emerged as two popular molecules that mimic viral and bacterial infections respectively and robustly activate the maternal immune system. In this study, we mimicked maternal viral infection by injecting poly I:C at various developmental time-points and followed the development of cortical GABAergic neurons in the offspring brains. Strikingly, in spite of an acute immune response, we revealed that the effect was both immediate, i.e. affecting proliferation of precursors of interneurons, and long-lasting, i.e. affecting migration of neuroblasts and maturation of cortical GABAergic neurons during late embryonic or early postnatal brain development.

One of our major findings is that the effect of maternal inflammation has a differential effect on GABAergic interneuron subtypes, highlighting subtype-specific vulnerability of neurons to maternally-derived stimuli. Hitherto, studies have revealed a convergent effect of genetic and environmental schizophrenia-related insults on PV+ interneurons^21,24^. However, while PV+ interneurons are crucial in synchronizing spike timing via gamma-oscillations, suppression of their activity alone has been found insufficient to reproduce schizophrenia-like phenotype in genetic or environmental mouse models of schizophrenia^68^. We show here a differential impact of maternal inflammation on interneuron subtypes and cortical regions that goes beyond PV+ interneurons. GABAergic interneurons in the somatosensory cortex were more affected in comparison to their counterparts in the motor cortex. Furthermore, the impact on PV+ interneurons was localized specifically to layer 4 and could be traced as early as P15, suggesting a specific developmental impairment. Importantly, we found a significant impact of maternal inflammation on SST+ interneurons, which were affected not only in layer 4 as PV+ interneurons but also in layer 2/3 despite both subtypes being derived from MGE-derived Nkx2.1+ progenitors. The overlap in layer 4 is of significance as SST+ interneurons in this layer target mainly PV+ fast-spiking interneurons unlike in layer 2/3 where pyramidal neurons are the primary targets^69^. Selective ablation of SST+ interneurons during development has been shown to arrest the maturation of PV+ interneurons in layer 5/6 suggesting an early role for SST+ interneurons in PV+ interneuron maturation^70^. Moreover, the decrease in MGE-derived interneuron populations might disrupt transient translaminar networks^71,72^ that modulate subsequent functional maturation of cortical circuits.

The effect of maternal inflammation at E9.5 on MGE-derived PV+ and SST+ interneurons in the offspring can be observed as early as E10.5, due to decreased proliferation of MGE-derived Nkx2.1+ progenitors. However, this effect was acute with proliferation returning to wild type levels by E14.5. This immediate and short-lived effect of maternal inflammation is in line with previous reports on cortical progenitors in an LPS model^27,73^. Contrary to the effect on MGE-derived subtypes, we found no change in CGE-derived VIP+ and CR+/SST− interneurons in animals exposed to maternal inflammation at E9.5, and neither CGE-derived COUP-TFII+ progenitors nor VIP+ and CR+/SST− interneurons in the mature cortex were affected. This is ostensibly due to a later ‘birthdate’ of CGE-derived interneurons between E12.5-E18.5^12^. However, even though the effect on CGE-derived COUP-TFII+ progenitors could be discerned by inducing inflammation at E12.5, we did not observe a significant decrease in VIP+ interneurons in somatosensory cortex at P15. This can be explained by the low relative production of VIP+ interneurons at E12.5 (approximately 5%)^12^ or that other but not somatosensory cortical areas are affected. In contrast, inducing maternal inflammation at E16.5 resulted in a marked increase in cycling MGE and CGE progenitors. Such variability of outcomes of maternal inflammation highlights selective vulnerability of interneuron progenitor pools that likely depends on expression of distinct transmembrane and intracellular signaling components at each stage of development.

In order to mitigate litter-to-litter variations due to maternal inflammation, recent guidelines have advocated for considering each litter as an experimental unit^74,75^. However, in this study we observed a greater within-litter than inter-litter variability. We thus sampled more than one offspring per litter (and analyzed at least 3 litters per condition). In addition, we used a linear mixed-model (LMM) analysis to appropriately represent this variation while considering the litter as a source of variation. This provided a better depiction of the range of effect of maternal inflammation.

It can hence be appreciated that the time of insult coupled with cellular or genetic vulnerabilities can produce varying outcomes on interneuron subtypes. Timing of the developmental insult will determine those developmental processes in the cortex that are affected, leading to variation in the cognitive phenotypes and severity of the affected offspring. Indeed, it has been shown that a difference in the timing of maternal inflammation leads to distinct behavioral dysfunctions^39,40^ (see also^76^ for review). Over the course of cortex development, maternal inflammation could interfere with: (1) interneuron precursor proliferation, (2) neuronal migration and positioning between cortical areas and within cortical layers, and (3) neuronal maturation and circuit connectivity. Our data shows that acute maternal inflammation affects multiple stages of interneuron development. In addition to the impact on precursor proliferation, described above, we show that migration of interneurons into the dorsal cortex was impaired as early as E14.5. The effect of maternal inflammation on the distribution of cortical GABAergic interneurons was maintained at E17.5 but could only be seen in the superficial cortical bins. This could be due to a greater proportion of GAD+ neuroblasts in maternal inflammation-exposed embryos migrating laterally and further work using time-lapse microscopy will be important to clarify the mechanism of this impairment. Likewise, while analysis of early postnatal stages showed a decreased density of GAD+ neuroblasts in regions of the developing cortex that correspond to both motor and somatosensory cortices, by P15 the effect persisted only in the somatosensory cortex. This suggests a differential regional impairment of cortical interneurons. Despite this, perinatal reduction in the number of GAD+ neuroblasts in the motor cortex might still affect the maturation of early cortical circuits and have functional outcomes that could not be measured in this study.

The effect of maternal inflammation on different stages of interneuron development (proliferation, migration and maturation) might be independent of one another or effects on neuronal migration and maturation might stem from the proliferation deficit. Although the major effect of maternal inflammation has been attributed to an acute increase in levels of pro-inflammatory molecules, the contribution of some low scale but significant chronic effects cannot be excluded. Furthermore, late embryonic inflammation can potentially affect the migration of neuroblasts directly and also those neuroblasts that start to mature in the cortical plate. Most probably, the effect of maternal inflammation on neuronal migration and maturation is complex and consists of direct and indirect components. Likewise, less pronounced differences in the number/distribution of PV+ and SST+ interneurons in adult offspring in comparison to early postnatal stage might be explained by a maturational delay of MGE-derived interneurons that stems from disturbed proliferation and direct chronic post-inflammatory effects.

Functionally, PV+ interneurons in our model showed a decreased maximum firing frequency in the absence of changes in input resistance. The maximum firing frequency in PV+ interneurons has been described to increase during postnatal development^77,78^, thus suggesting a maturational delay of PV+ interneurons in our study due to maternal inflammation. This maturational delay is also corroborated by decrease in dendritic complexity of maternal inflammation-affected PV+ interneurons. Alterations in cellular properties of GABAergic interneurons can alter the perisomatic inhibition of pyramidal neurons thus altering the excitation-inhibition balance in the neocortex^78^. Furthermore, human studies show widespread changes in neonatal cortical connectivity due to increase in the level of maternal proinflammatory cytokines^79,80^. Importantly, GABAergic interneurons are necessary for maturation of early cortical circuits that provide the foundation for adult cortical connectivity^71,72^. Thus, maturational delay of GABAergic interneurons due to maternal inflammation can have broad effect on cognitive properties of the affected offspring. In particular, this might lead to reduced cortical plasticity and in turn impaired learning and memory in the older mice.

While this study focuses on GABAergic interneurons, maternal inflammation can also have an effect on populations of pyramidal neurons in the cortex ^27,81^. Furthermore, thalamic and subplate neurons are also likely to be affected at early gestational stages since they are born between E10-E12^82,83^. A disruption of either of these neuronal populations could impact the maturation of GABAergic interneurons and hamper their functional integration into cortical circuits^84,85^, thus adding to the overall effect of maternal inflammation on development of GABAergic interneurons.

The poly I:C model of maternal inflammation presents high construct validity and parallels the effects seen in humans^3,76^. Based on translation of brain development milestones, the early time-point of induction (E9.5) overlaps with the first trimester of human pregnancy^47^ as well as with the onset of neurogenesis of cortical GABAergic interneurons^86^. Given the significant evidence for impairment of cortical GABAergic interneurons in human postmortem brain studies of patients with mental illness^87–89^, the dysfunction of developmental programs for GABAergic interneurons might have causative effect on cognitive phenotype of the patients. Further studies of GABAergic circuitry development in environmental and genetic mouse models mimicking human developmental insults will reveal commonalities and differences in the affected subtypes of GABAergic interneurons across all developmental processes. Such data will provide mechanistic insight into the etiology of human neurodevelopmental disorders and will identify therapeutic windows during brain development when impairments in GABAergic circuitry can be corrected.

## Acknowledgements

We are grateful to Prof. Yuchio Yanagawa (Gunma University, Japan) for sharing the GAD67-EGFP knock-in mouse line. We also wish to thank Dr. Yasuko Antoku for assistance with microscopy, Johanna Meichsner (University of Mainz) for help with statistics, and members of the Khodosevich lab for constructive discussions relating to this study.

The work was supported by Novo Nordisk Hallas-Møller Investigator grant (NNF16OC0019920), Lundbeck-NIH Brain Initiative grant (2017-2241) and DFF-Forskningsprojekt1 (8020-00083B) to KK and German Research Foundation (DFG) grant within the Collaborative Research Center (SFB) 1080 “Molecular and Cellular Mechanisms of Neural Homoeostasis” to JvE.

## Author Contributions

This study was conceived by NAV, MPN, UP, and KK. NAV and MPN performed the poly I:C injections and downstream experiments including design, collection, analysis and interpretation of data. JG, DW, SM, DGG collected and analyzed specific developmental stages. UPF performed and analyzed IL-6 ELISA. IK performed and analyzed the western-blotting for PV. MKM and JvE performed and analyzed electrophysiology. KSH contributed to the design, execution and analysis of behavioral experiments. NAV and KK jointly wrote the paper. All authors read and approved the manuscript.

## Conflict of Interest Statement

The authors declare that they have no potential conflicts of interest

Supplementary information is available at MP’s website.

## Notes

#### Summary of Updates

Following peer review, significant changes were made to the manuscript including text and figure changes. New supplementary figures have been added to support the findings and have been made to the original figures.

