## Supplementary Figure Legends for "Maternal inflammation has a profound effect on cortical interneuron development in a stage and subtype-specific manner"

**Supplementary Material**

**Supplementary Figure Legends**

**Supplementary Figure 1: Exposure to maternal inflammation affects both sexes of offspring**

(a) PPI was found to be significant only at 72dB but not at the other pre-pulse frequencies.

(b,c) PPI was found to be reduced in both females and males separately at 72dB but did not reach statistical significance.

(d-f) Both females and males showed a reduction in time spent in the social zone but did not reach statistical significance (c). On the other hand, time spent in social zone in 1-minute bins was found to be significantly reduced for both males and females exposed to maternal inflammation.

(g,h) Females (g) but not males (h) showed an increase in total distance travelled in the open-field test.

(I,j) Velocity of running differed in both females (i) and males (j) though the effect was greater in females.

Comparison of means by 2-way ANOVA to analyze prepulse inhibition, with prepulse as the repeated measure in (a-c), Multiple t-test with Holm-Sidak correction in (d,g,h) and 2-way ANOVA with Sidak’s multiple comparison test in (e,f,i,j) (mean±SEM are shown, p-values denoted as asterisks with <0.05 shown as * and <0.005 shown as **).

**Supplementary Figure 2: Maternal inflammation does not affect distribution of neuroblasts at the pallial-subpallial boundary and cortical thickness in the developing brain**

(a) No evidence for arrest of migrating EGFP+ neuroblasts at the pallial-subpallial boundary was observed in poly I:C exposed embryos at E14.5.
(b) Total cortical thickness (WM-MZ), cortical plate (CP) and subventricular zone (SVZ) thickness of the developing cortex are not affected in poly I:C exposed embryos at E14.5.
(c) Marginal zone (MZ), cortical plate (CP) and intermediate zone (IZ) thickness is not affected

in poly I:C exposed embryos at E17.5.

Comparison of means by Student’s t-test with Holm-Sidak’s correction in (b,c). Scale bar: 50 µm.

**Supplementary Figure 3: Relative distribution of EGFP+ neuroblasts in the developing cortex at P3, P6 and P9**

(a-c) Representative images of P3 (a), P6 (b) and P9 (c) somatosensory cortex showing the distribution of EGFP+ neuroblasts in PBS and poly I:C exposed animals.

(d,e) Graphs showing comparable proportions (in percentage) of EGFP+ neuroblasts in PBS- and maternal inflammation-exposed offspring across the cortex at P3 (a) and P6 (b).

Comparison of means by 2-way ANOVA with Sidak’s multiple comparisons in (d,e). Scale bars: 50 µm.

**Supplementary Figure 4: Maternal inflammation-exposed offspring do not show an increase in apoptosis of cortical GABAergic interneurons during development**

(a) Representative images demonstrating labeling of apoptotic GABAergic neuroblasts by double-staining for activated caspase-3 and EGFP.
(b) Exposure to poly I:C during early development does not increase the number of activated caspase-3+ or activated caspase-3+/EGFP+ cells at E17.5, P3 or P6.

Comparison of means by Multiple t-test with Holm-Sidak correction in (b). Scale bar: 50 µm.

**Supplementary Figure 5: Interneuron subtype-specific effects of maternal inflammation**

(a) Anatomical region considered as the medial prefrontal cortex (mPFC): secondary motor cortex (M2), cingulate (Cg) and prelimbic (PrL) cortices.

(b,c) Representative images from PBS and poly I:C exposed animals at P15 (b) and graphs (c) showing lack of changes in PV+ or SST+ cell numbers in the mPFC.

(c,d) Maternal inflammation at E9.5 does not affect the distribution of the CGE-derived VIP+ (c) or CR+/SST- (d) interneuron subtypes at P15.

(f-h) Representative images from PBS and poly I:C exposed animals at P15 (f) and graphs (g,h) demonstrating effect of maternal inflammation at E12.5 on CGE-derived VIP+ (g) or CR+/SST- (h) interneurons. Note that L5 VIP+ interneurons in (g) are virtually absent in maternal inflammation-affected offspring.

(i,j) Maternal inflammation at E12.5 does not affect the distribution of the MGE-derived PV+ (i) or SST+ (j) interneuron subtypes at P15.

Comparison of means by Student’s t-test with Holm-Sidak correction for multiple comparisons in (c,d,e,g-j) (mean±SEM are shown). Scale bars: 50 µm.

**Supplementary Figure 6: Western blotting of proteins from P60 brains shows a decrease in levels of parvalbumin**

(a) Representative western blots showing expression of parvalbumin (PV) and loading control GAPDH in cortices of P60 PBS and poly I:C exposed mice.
(b) Estimation of PV levels shows a decrease in poly I:C exposed mice.

Comparison of means by Student’s t-test (mean±SEM are shown).

**Supplementary Figure 7: Ventricular zone proliferation is not affected by maternal inflammation**

(a,b) Ventricular zone (VZ) proliferation is not affected at E10.5 in MGE (a) and at E14.5 in CGE (b) respectively.

Comparison of means by Student’s t-test (mean±SEM are shown).

**Supplementary Figure 8: Lack of poly I:C lot-specific effects**

(a-f) Absence of lot-specific effects. Panels (a,b), (c,d) and (e,f) correspond to Figure 3e,g, Figure 4c,d and Figure 4e,f respectively. Closed and Open symbols represent separate lots.

Note that only those experiments are shown where significant differences were found between PBS- and poly I:C-exposed offspring. Statistical comparisons are shown in Suppl. Table 1.

**Supplementary Figure 9: Lack of litter-specific effects for PBS and poly I:C injections**

(a-k) Absence of litter-specific effects. Panels (a-d) correspond to Figure 6b,g,i,j , panels (e-g) correspond to Figure 3a-c, panels (h,i) correspond to Figure 3e,g and panels (j,k) correspond to Figure 4c,d respectively. Closed, Open and Closed coloured symbols represent animals taken from different litters. Note that only those experiments are shown, where significant differences between PBS- and poly I:C-exposed offspring were found. Statistical comparisons are shown in Suppl. Table 4.

**Supplementary Tables**

**Supplementary Table 1: Lack of Poly I:C lot-specific differences in the maternal inflammation-affected offspring**

**Supplementary Table 2: Litter sizes and Abortion rates in PBS and Poly I:C injected dams**

**Supplementary Table 3: Number of animals included for each analysis**

**Supplementary Table 4: Lack of litter-specific differences for PBS- or Poly I:C-affected offspring**

**Supplementary Table 5: Confirmation of significant differences between PBS and Poly I:C-affected mice by Linear Mixed Model statistics**
